# Genome analysis and data sharing informs timing of molecular events in pancreatic neuroendocrine tumour

**DOI:** 10.1101/232850

**Authors:** Rene Quevedo, Anna Spreafico, Jeff Bruce, Arnavaz Danesh, Amanda Giesler, Youstina Hanna, Cherry Have, Tiantian Li, S.Y. Cindy Yang, Tong Zhang, Sylvia L. Asa, Benjamin Haibe-Kains, Suzanne Kamel-Reid, Monika Krzyzanowska, Adam Smith, Simron Singh, Lillian L. Siu, Trevor J. Pugh

## Abstract

Neuroendocrine tumours (NETs) are rare, slow growing cancers that present in a diversity of tissues. To understand molecular underpinnings of gastrointestinal (GINET) and pancreatic NETs (PNETs), we profiled 45 tumours combining exome, RNA, and shallow whole genome sequencing, as well as fluorescent in situ hybridization. In addition to expected somatic mutations and copy number alterations, we found that PNETs contained a highly consistent copy neutral loss-of-heterozygosity (CN-LOH) profile affecting over half of the genome; a greater percentage than any cancer analyzed to date. Our data indicates that onset of extreme autozygosity may be progressive, associated with metastasis, and initially triggered by the loss of *DAXX/ATRX*, and subsequent biallelic loss of *MEN1*. We confirmed this molecular timing model using targeted clinical sequencing data from an additional 43 NETs made available by the AACR GENIE project. Against this background of CN-LOH, several chromosomal regions consistently retained heterozygosity, suggesting selection for crucial allele-specific components specific to PNET progression and potential new therapeutic targets.

**Statement of significance:** We have discovered that pancreatic neuroendocrine tumours contain a characteristic pattern of copy neutral loss-of-heterozygosity affecting the majority of the genome following mutations of *MEN1* and *ATRX/DAXX*. Against this background of loss-of-heterozygosity, specific genomic regions are consistently retained and may therefore contain vulnerable therapeutic targets for pancreatic neuroendocrine tumours.

## Introduction

Neuroendocrine tumours (NETs) are cancers of peptide hormone-producing neuroendocrine cells in glands such as pituitary and parathyroids, and scattered throughout many organs including the thyroid, thymus, lung, pancreas, and gastrointestinal tract. NETs of the gasteroenteropancreatic system are increasing in incidence faster than almost any other cancer, growing from an incidence of 1.1 to 5.3 per 100,000 individuals between 1973 and 2004^1^. Both pancreatic neuroendocrine tumours (PNETs) and gastrointestinal neuroendocrine tumours (GINETs) often present with local and distant metastases at diagnosis, limiting the efficacy of standard medical treatments^1^. Each present with different genetic profiles and clinical development, however they are also often managed as a group and have some overlap in response to mTOR-targeted therapy^2,3^.

Both pancreatic and gastrointestinal NETs harbour their own distinct genomic landscapes, each with few recurrent mutated genes. PNETs are characterized by mutations in chromatin modifiers *MEN1, DAXX* and *ATRX*, while GINETs show recurring loss-of-function mutations in *CDKN1B*^1,4-7^. The loss of *DAXX* or *ATRX* in PNETs is associated with an alternative lengthening telomere (ALT) phenotype that represents a more aggressive subtype and is prone to chromosomal instability^8^. Additionally, there is a biologically distinct subgroup of metastatic PNETs that is enriched for mutations in both *ATRX/DAXX* and *MEN1^4^*. This subgroup not only has longer overall survival^4^, but the concomitant mutation of both sets of genes suggests a compensatory mechanism that favours tumour stability over *ATRX/DAXX* mutations alone. Furthermore, both PNETs and GINETs contain heterogenous karyotypes in which haploinsufficiency through loss-of-heterozygosity (LOH) is a suggested mechanism for tumour progression^9,10^.

In this study, we report a comparative study of genome alterations between PNETs and GINETs. Unexpectedly, we uncovered a highly recurrent pattern of copy neutral loss-of-heterozygosity that is distinct to metastatic PNETs with ATRX/DAXX and MEN1 mutations (MAD+) in our discovery cohort. To further examine the consistency of this LOH signature, we extended our analysis to include a validation cohort, publicly available array CGH datasets, and targeted-panel sequencing from the American Association for Cancer Research (AACR) Project Genomics Evidence Neoplasia Information Exchange (GENIE) dataset. We uncovered a trend that places MAD+ mutations as an initiating molecular event that precedes the characteristic LOH pattern and genome doubling in PNETs. Tumour with this karyotypic signature were associated with metastasis and a more aggressive clinical course. Herein, we propose a model for the highly conserved progression of molecular events that defines an aggressive subtype of PNETs.

## Results

### Characterization of karyotypes

To compare allele-specific copy-number (ASCN) alterations shared by GINETs and PNETs, we profiled 45 NETs combining exome sequencing (2 PNET metastases, and 5 primary/metastasis pairs from 2 PNETs and 3 GINETS), shallow whole genome sequencing (13 PNETs, 10 GINETs), and fluorescent in situ hybridization (5 PNETs, 7 GINETs) (Supp. Tables 1 and 2). In addition to expected somatic mutations and copy number alterations (CNAs), we found that >50% of the genome was subject to recurrent chromosomal loss-of-heterozygosity (LOH) distinct to PNETs, affecting a greater fraction of the genome than any tumour type analyzed to date (Fig. 1). Against this background of extreme autozygosity, seven chromosome arms retained heterozygosity in all tumours, suggesting selection for retention of allele-specific gene expression within these loci for PNET progression. To rule out the possibililty of inherited germline LOH, we analyzed patient-matched white blood cells and found all patients to have normal diploid heterozygous genomes (Supp. Fig. 1a).

**Figure 1.**
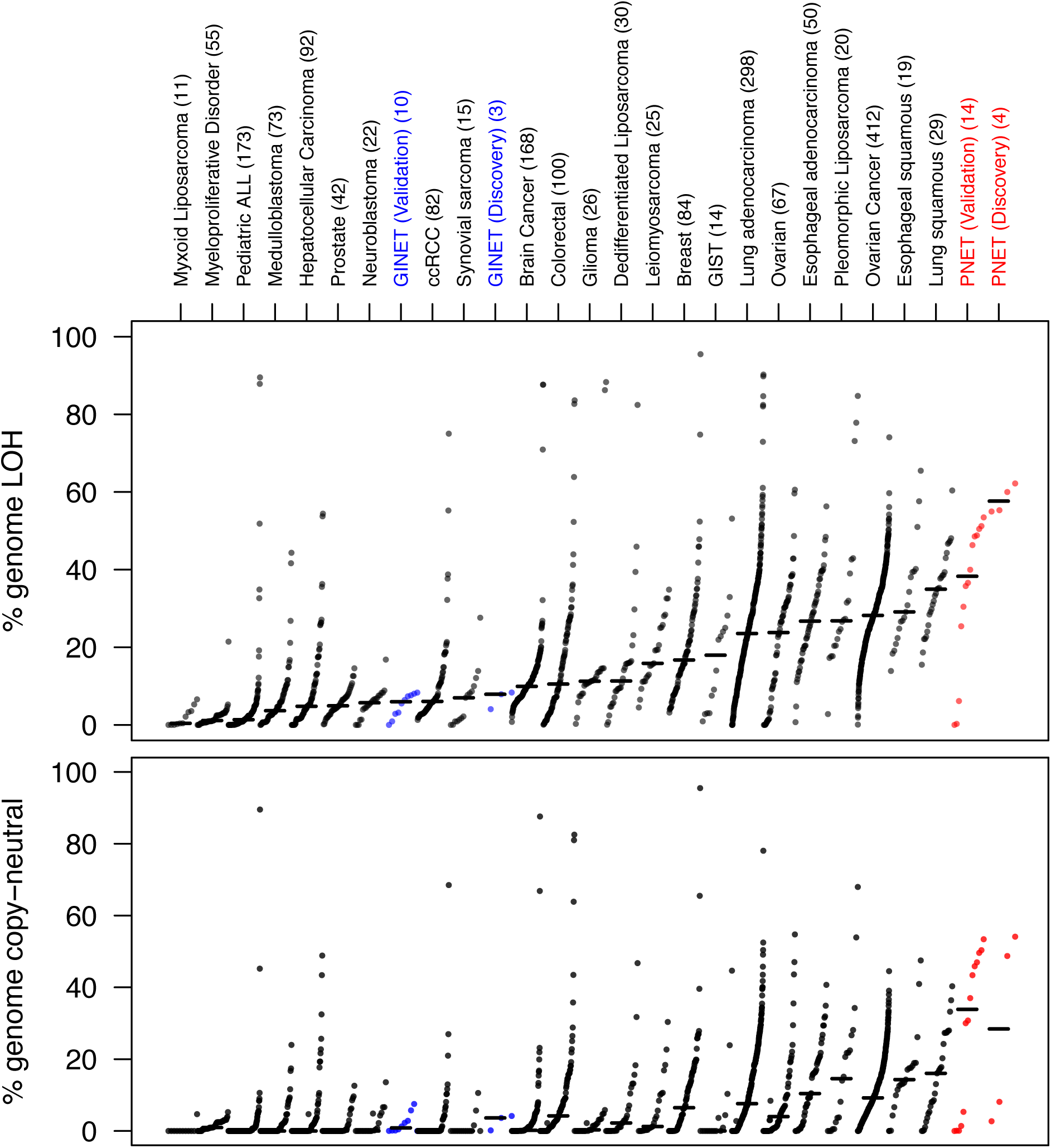
The percentage of the genome that is LOH plotted for a subset of the TCGA pan-cancer analyzed tumours. PNETs (red) and GINETs (blue) from the discovery and validation cohort are plotted in reference to all other tumour types. The top track indicates the percentage of the total genome that is LOH, while the lower tracks represents the percentage of the total genome that is copy-neutral.

As an initial discovery set, we generated allele-specific copy-number profiles using exome sequence analysis of 14 NETs primary/metastasis pairs from 7 patients. Overall, GINETs had few CNAs beyond characteristic loss of chromosome 18 in 4 of 6 tumours from 2 of 3 cases and ~2.6% of the genome had LOH on average. In contrast, the 8 PNETs from 4 patients showed a recurrent signature of gains and losses consistent with previous CGH studies including gains in chromosomes 4, 5, 12, 14, 17, 19 and 20^11^^-^^13^ (Supp. Fig. 1b). Moreover, in all PNET samples, we found a distinct pattern of copy-neutral loss of heterozygosity (CN-LOH) that affected over half of the genome in each tumour, yet consistently preserved heterozygosity of 4q, 5p, 7q, 14, 17, 19, and 20q (Table 1). Such a pattern has not yet been observed in any other tumour type and we therefore sought to confirm this pattern using additional genomic platforms.

**Table 1.**
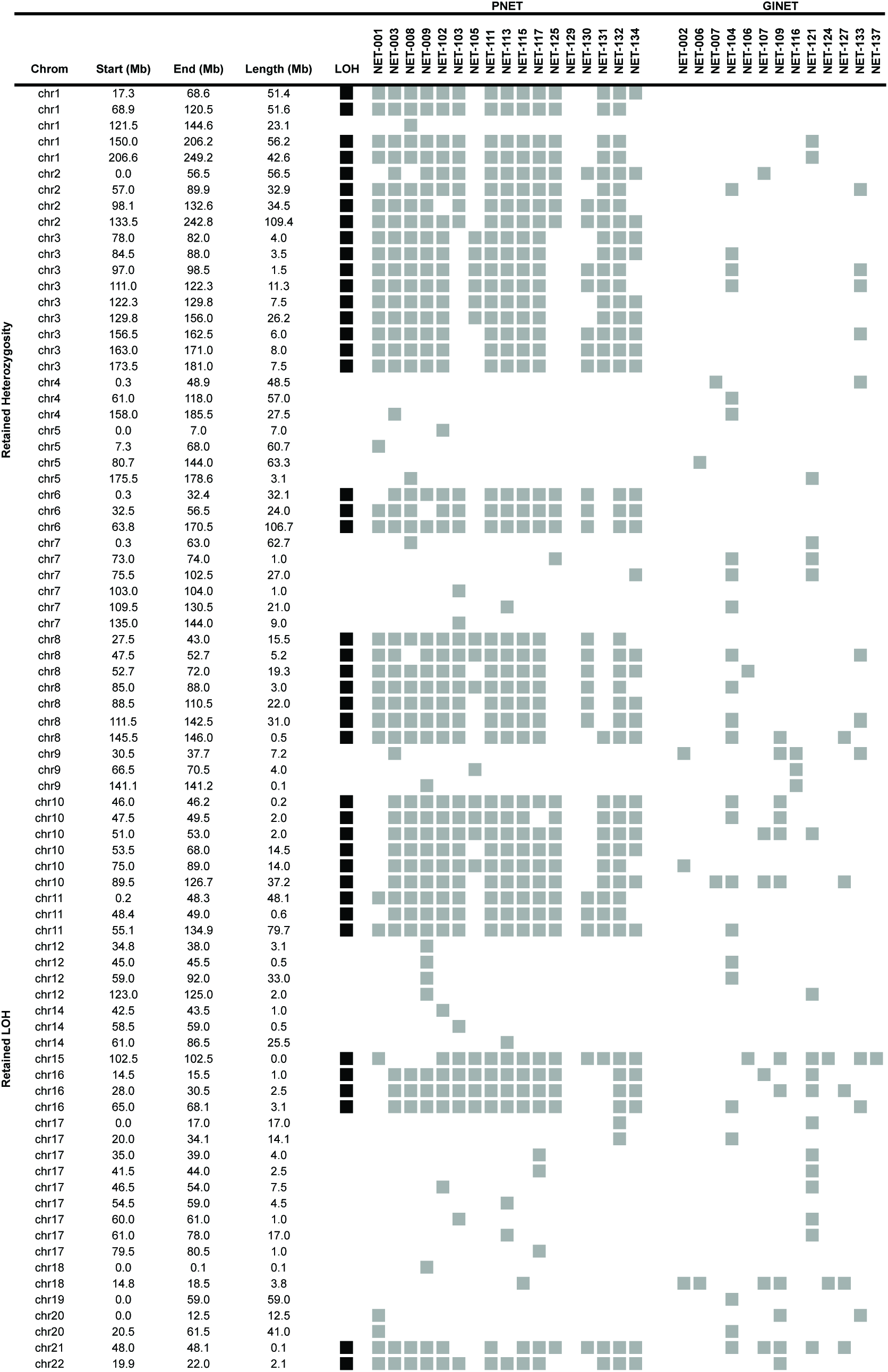
Genomic coordinates based on the hg19 reference genome for consensus heterozygous and LOH regions in PNETs within the discovery and validation cohort. A heterozygous region is defined as a single LOH event in one sample across all 17 PNETs, while a LOH region is defined as 13+ samples harbouring an LOH at that given region.

To confirm this PNET-specific CN-LOH pattern, we re-analyzed the DNA from our discovery cohort using the Affymetrix SNP6 platform as well as an analysis of expressed germline variation using matched RNA sequencing (RNA-seq) (Supp. Fig. 1c). Copy number alterations and CN-LOH regions were highly concordant with those inferred from exome analysis, with segmental overlap ranging from 92 to 100% (Supp. Fig. 2a). Similarly, LOH-segments inferred from RNA-Seq data showed a high degree of overlap with the regions identified from WES (84-88%) (Supp. Fig. 2b). We attribute the small, incomplete overlap of DNA/RNA segements to the presence of normal cell admixture in tumour tissues and potential for variable gene expression levels across individual cancer. Overall, the regions of CN-LOH and retained heterozygosity in PNETs were cross-validated by strong concordance between DNA and RNA sequencing approaches.

### Validation of the copy-neutral LOH signature

To determine the frequency of the aneuploidy patterns seen in our discovery cohort, we performed shallow whole genome sequence (sWGS) analysis (Supp. Fig. 3) of an extended 23-sample validation cohort comprised of 13 PNETs (6 metastatic, 7 non-metastatic) and 10 GINETs (9 metastatic, 1 unknown) (Supp. Table 2.). As a complementary approach and to validate the copy-status of these samples, we used FISH analysis with centromeric probes targeting chromosomes 3, 7, 10, 17 and 18 on 12 of the 23 samples analyzed with sWGS (Supp. Table 3). While both FISH and sWGS were concordant in 9/12 samples for all chromosomes probed, there were some discrepancies that may have resulted from differences in clonal heterogeneity within the tumour (Supp. Table 4). Of these discrepant samples, NET-102 and −134 were discordant in one chromosome each, with raw sWGS coverage and zygosity data clearly supporting different copy states from those seen from FISH counts. Chromosome counts from FISH and sWGS were discrepant at 2/5 loci in NET-116, suggesting PNETs may contain a mixture of subclones at different points along the development of the characteristic CN-LOH signature. Overall, consistent with our discovery cohort, the validation PNETs exhibited significantly more LOH than GINETs in chromosomes 1, 2, 3, 6, 8, 10, 11, 16, and 22, while chromosomes 15, and 21 were more likely to be found in a heterozygous state (Fig. 2).

**Figure 2.**
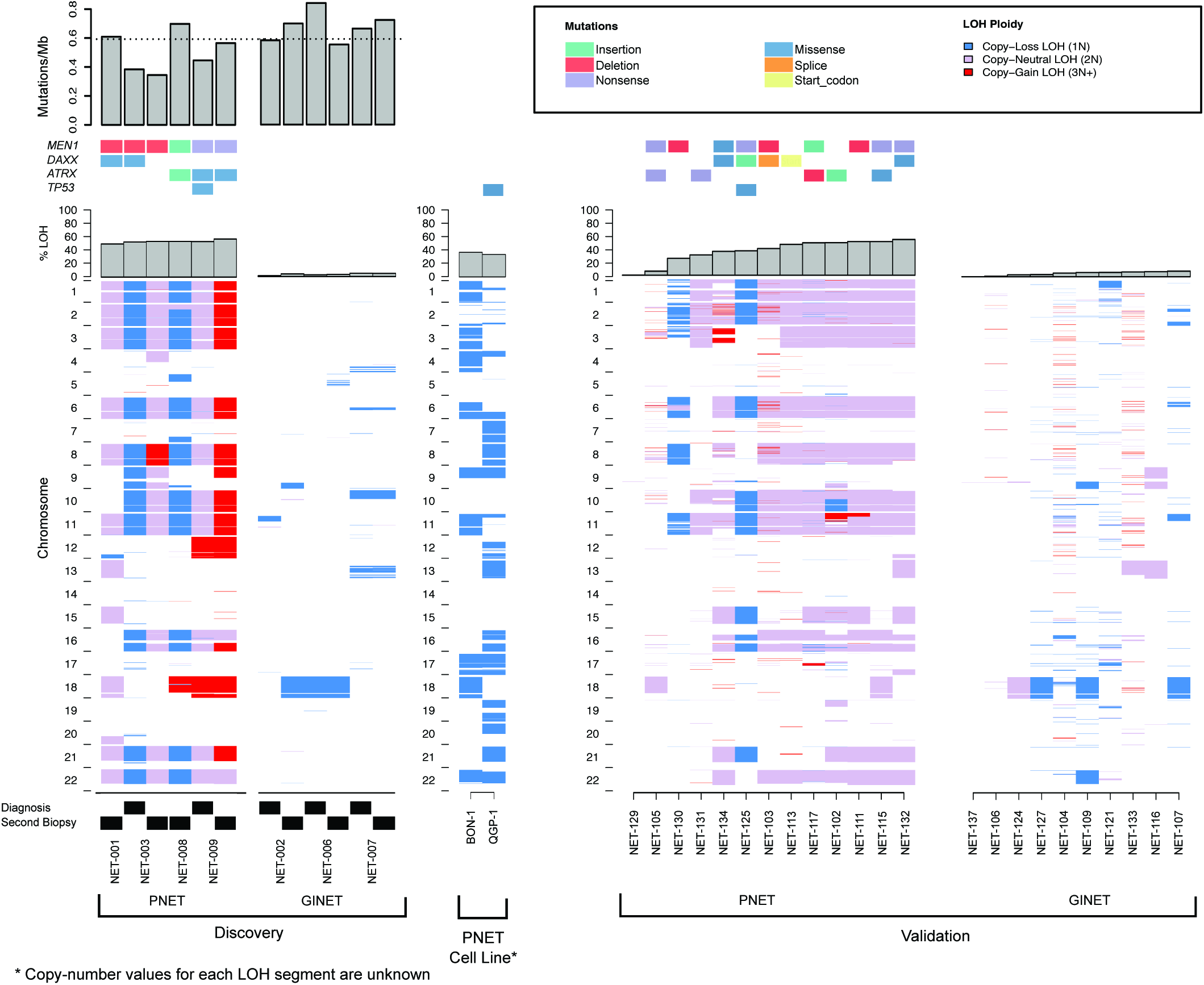
Loss-of-heterozygosity profiles depicted as being either copy-loss/haploid (blue), copy-neutral/diploid (purple), or copy-gain/triploid+ (red) for each sample in the discovery, validation, and PNET cell line data. Mutations on characteristic NET genes are indicated above their corresponding sample, as well as the mutational burden of the sample.

To refine regions of consistent zygosity in PNETs, we explicitly defined highly conserved LOH and heterozygous regions common across discovery and validation cohorts. We defined a conserved heterozygous region as a heterozygous region present in 16/17 samples while conserved LOH regions required 13/17 samples to show evidence of LOH. More relaxed requirements were used for LOH to compensate higher variability for LOH estimations in tumour tissues of low cancer cell content (Table 1). Metastatic PNETs showed a stronger trend towards conserved LOH regions compared to non-metastatic PNETs (OR 1.73, 95% CI [0.94, 3.19]; p=0.079) (Supp. Table 5). Of the conserved LOH regions, chromosomes 1, 10, 16 and 18 were strongly associated with metastasis, while LOH of chromosomes 6, 8, 21 and 22 were weakly associated with metastasis (Supp. Table 5). While statistical significance was not achieved, we believe that this association may be obscured by the small sample size of metastatic (n=6) and non-metastatic PNETs (n=8) in our validation cohort. Thus, a potential link may exist between disease progression and chromosomal instability in PNETs that may be tested in larger populations.

To evaluate whether this CN-LOH signature is recapitulated in PNET model organisms, we evaluated genome-wide zygosity using publicly available exome sequence data from two well-characterized metastatic PNET cancer cell lines, BON-1 and QGP-1^14^. A reanalysis of this dataset corroborated previous reports of non-pathogenic germline variants *MEN1* p.T541A and *ATRX* p.E929Q in BON-1, and *MEN1* p.T541A and *ATRX* p.F847S in QGP-1. Consistent with our discovery cohort, both PNET cell lines exhibited a similar near-global LOH signature (Supp. Fig. 4), with the BON-1 cell line displaying a more concordant pattern of LOH to our PNET samples (Fig. 2). This may be a result of BON-1 and QGP-1 representing different subtypes or stages of PNET, with BON-1 being more representative of the metastatic PNETs we analyzed^15^.

### Extended validation of CN-LOH signature

Given the significance of highly-recurrent CN-LOH apparent in PNETs, we sought to compare the magnitude of this signature with a subset of The Cancer Genome Atlas pan-cancer data set with available allelic-specific copy number profiles^16^. Across 1,944 tumors from 27 cancer types, the median percentage of the genome affected by LOH was 14%, with lung squamous cell carcinoma having the largest percentage at 35%. At 54% genome-wide LOH, metastatic PNETs from our discovery cohort were an extreme outlier compared to all other tumor types in the pan-cancer cohort (Fig. 2). Conversely, GINETs had only 4% of the genome affected by LOH. LOH profiles derived from the validation cohort analyzed by less-sensitive sWGS were still greater than the pan-cancer cohort with a LOH fraction of 38%, 11 of which were copy-neutral. Thus, metastatic PNETs harboured the largest genomic fraction of LOH of any tumour analyzed in this panel (p<5.1x10^−7^; one-sided t-test) with a strong trend towards copy-neutrality.

While comparative genomic hybridization (CGH) is unable to detect CN-LOH, published studies using this technique have uncovered a wide diversity of CNAs in NETs^11-13,17-22^. To compare these historical NET copy number data with our own, we performed a meta-analysis of 226 NETs from 8 previous reports (Supp. Table 6-14). By clustering the total-allelic copy number (TCN) profiles of our NETs and published datasets (Online methods), we were able to show that 7 clusters clearly divided into groups of high and low aneuploidy in addition to clustering minor variations within each group (Supp. Fig. 5). PNETs in our discovery and validation cohort largely grouped together in clusters 1 and 5 while showing the same overall trend of gains on chromosomes 4, 5, 7, 9, 12, 13, 14, 17, 18, 19, 20. Clusters 3 and 7 displayed an inverse of the karyotypes observed in clusters 1 and 5, composing largely of haploid karyotypes that mimicked those seen in NET-008, −121, −125, and −130 of our discovery and validation cohorts. It is possible that this cluster represents a subset of NETs that have yet to undergo genome doubling as copy-loss frequently occurs as a precursor event^23^. Cluster 4 was comprised of a small cohort of NETs that were defined by only chromosome 18 loss, a feature well characterized in small intestine NETs^5,7^. Similarly, GINET samples in both cohorts were localized to cluster 2 which was characterized by an absence-of or minimal copy-number aberrations, featuring predominantly a diploid karyotype. Finally, cluster 6 lacked any defining features other than gains of chromosomes 4, 7, and 17 in a few samples. Two PNET samples in our validation cohort were found within this cluster, however, attributing a salient feature was difficult as both samples yielded poor signal and low-confidence copy-states.

Based on our previous observation that LOH status in PNETs was associated with metastatic status, we next sought to identify whether chromosomal instability can be used as an indirect measure of the LOH signature. By ranking the fraction of genome that was chromosomally unstable, we were able to separate the copy-number profiles into high-chromosomal instability (high-CI) and low-chromosomal instability (low-CI) groups based on the bimodal nature of the nominated clusters (Supp. Fig. 5b). NETs with high-CI were more likely to be metastatic (OR 2.39, 95% CI [0.97, 5.85]; p=0.057; I^2^=25.20) (Table 2) and the majority of the high-CI NETs were those found in clusters 1, 3, 5, 6, and 7 (Supp. Table 15-21); all tumours that largely follow the proposed karyotype associated with the CN-LOH signature observed in our discovery and validation cohort. Analyzing each cluster independently, clusters 1 and 5 had the strongest trend towards being metastatic (OR 2.57, 95% CI [0.49, 13.52] and OR 2.63, 95% CI [0.45, 15.39] respectively) as well as cluster 5 favoring high-CI PNETs over GINETs (OR 8.16, 95% CI [1.44, 46.32]). Thus, we suggest that using this karyotype as an indirect measure of the copy-neutral LOH signature is a marker for late-stage neuroendocrine tumors and the development of metastasis.

**Table 2.**
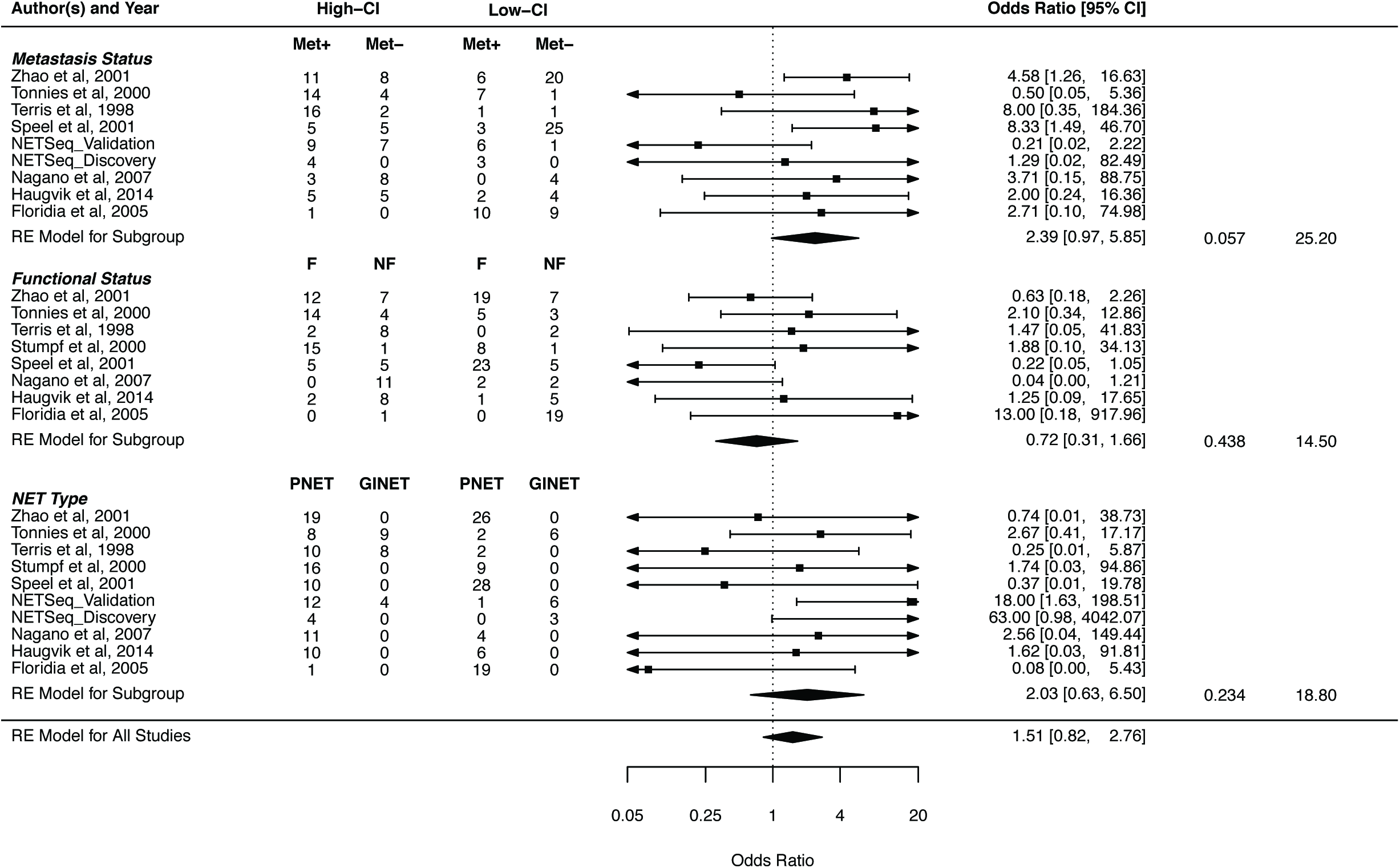
A meta-analysis of the CGH datasets for the highly aneuploid NET tumours (High-CI) against the low aneuploid NET tumours (Low-CI). The parameters being compared are the metastasis status of the tumour type (Met+: Metastasis present, Met-: No metastasis), the functional status (F: functional, NF: non-functional), and the NET type (PNET: pancreatic neuroendocrine tumour, GINET: gastrointestinal neuroendocrine tumour).

### Alternative Lengthening of Telomere phenotype

To test whether the loss of *DAXX/ATRX* and *MEN1* was associated with an ALT phenotype as previously reported^4,8^, we compared the lengths of telomeres between 13 PNETs and 10 GINETs with sWGS data. We observed more variable and significantly longer telomere lengths in PNETs compared to GINETs, suggestive of an ALT phenotype (Supp. Fig. 9). With the exception of NET-132, all PNETs had longer telomeres than GINETs; while NET-132 was only marginally shorter than the longest GINET telomeres. These results confirm reported associations, where high-CI status and MAD+ status are associated with ALT in PNETs. As such, it is likely that the chromosomal instability described in previous studies is actually this well-conserved CN-LOH karyotype.

### Molecular timing of PNET progression

True to the nature of NETs, the tumours analyzed by exome sequencing had low mutation rates (0.59 mutations / coding Mb) and few recurrently mutated genes (Supp. Table 22). With one exception, mutations in coding regions were proportionally scattered between LOH and heterozygous segments for each sample indicating no consistent enrichment of either regions (Supp. Fig. 6).

All 4 PNETs in our discovery cohort were MAD+ in the primary tumour. While mutations in these genes were found in both primary and metastatic tissues of three patients, a fourth patient (NET-003) lost a mutated *DAXX* allele through copy number loss found only in the metastasis. Similarly, patient NET-009 had a metastasis-specific loss of a *TP53* missense mutation, futher illustrating ongoing chromosomal instabililty between primary and metastatic tissues (Fig. 2).

To establish the mutational profile of all the samples within the discovery and validation cohort, we applied a deep-sequencing panel targeting 21 recurrently mutated genes in PNETs reported by Scarpa and colleagues^24^ (Supp. Table 23). This panel recapitulated all mutations detected by exome sequencing of the discovery cohort. Within the validation cohort, 15/16 PNETs harbored a mutation in at least one of the three genes; 10 of which had mutations in both *MEN1* and *ATRX* or *DAXX* (MAD+) (Fig. 2). The remaining PNET sample did not have either of these mutations, nor harbor the CN-LOH signature seen across all other samples, implicating a possible association between MAD+ and CN-LOH signature. As expected, all GINETs were MAD-.

To establish the chronological order of molecular events in PNETs, we mapped the allele fractions of mutations in *MEN1*, *ATRX* /*DAXX*, and other genes from the 21-gene panel relative to, and on the background of, mis-segregation events (Supp. Fig. 7). After correcting for tumour purity, we found cancer cell fractions of MAD mutations were consistent with presence on all chromosomal copies within each tumour, suggesting these mutations occur prior to global CN-LOH (Fig. 3). One case contained loss of a *DAXX* mutation in a metastatic sample, suggesting that this mutation may no longer be necessary for PNET development following occurrence of an unstable karyotype.

To investigate the association and molecular timing of MAD mutations and CN-LOH profile in PNETs, we leveraged clinical panel sequencing data from PNETs included in the initial release of the American Association for Cancer Research (AACR) Project Genomic Evidence Neoplasia Information Exchange (GENIE) dataset^25^. Of the 43 samples that contain both copy number and somatic mutation data in the GENIE v1.0 data freeze, 29 were MAD+ and 14 were MAD-. Of the MAD+ population, all samples display the characteristic CN-LOH signature while only 1 of the 14 MAD-samples showed these alterations (MAD+) (Fig. 4). While 20 of the 29 MAD+ samples contained both *MEN1* and *DAXX/ATRX* mutations, we hypothesize the remaining 9 tumours may harbour alternate mutational or epigenetic alterations not detected by our targeted sequencing approach.

**Figure 3.**
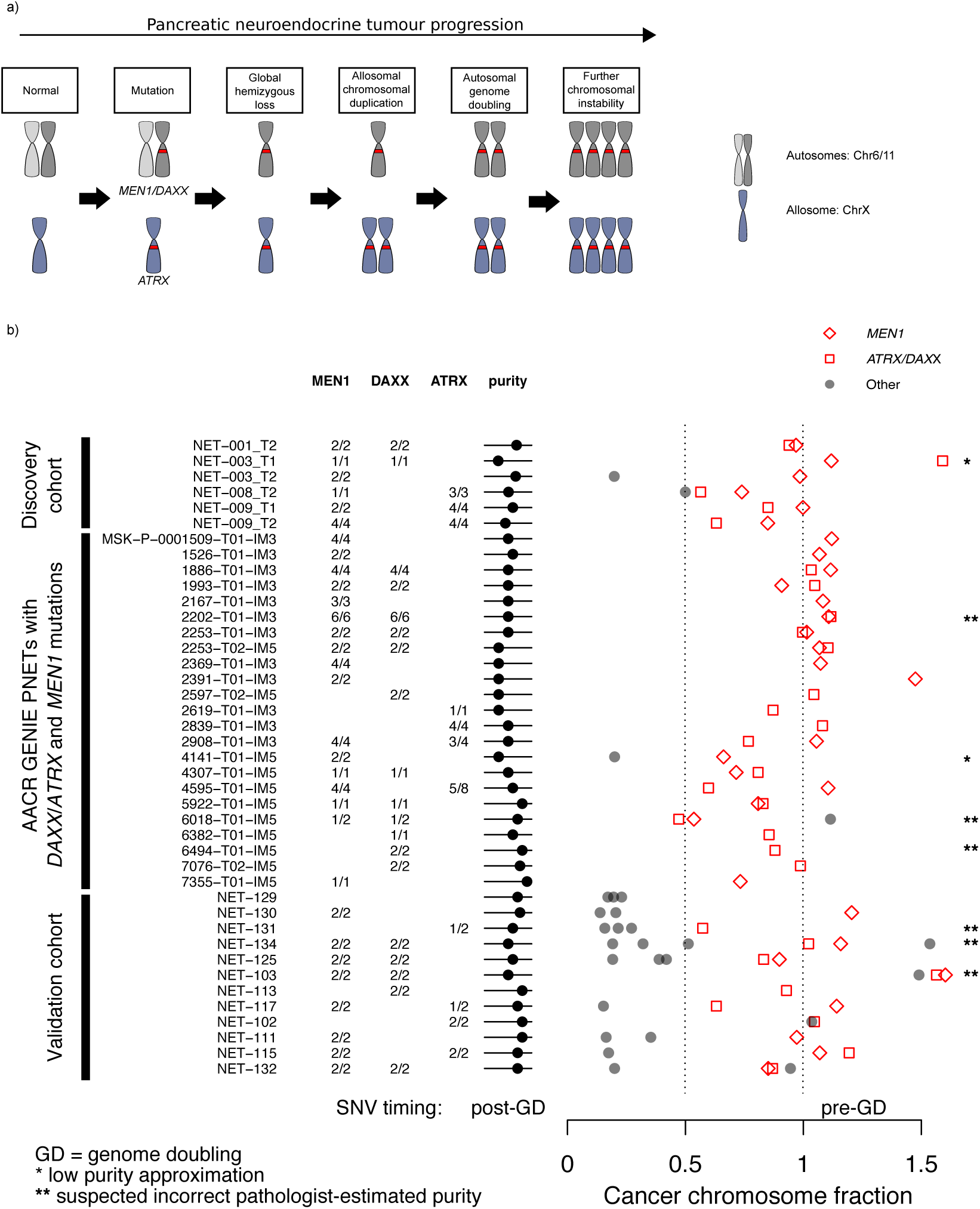
**a)** Proposed timeline of mutational events. Acquisition of *MEN1* and *DAXX* mutations are represented as a red band on the grey autosomal chromosomes, while *ATRX* is shown on the blue allosomal chromosome. **b)** Estimations of the theoretical tumour allelic fraction for *MEN1* (red diamond), *DAXX/ATRX* (red square), and other gene-level mutations (grey circles) for the copy number model (Number of ALT alelles/Ploidy) that best represents the pathologist-estimated purities across the different cohorts. A fraction of 1.0 indicates a homozygous variant, and 0.5 a heterozygous variant. Any deviations from these values represent variance in the observed allelic fractions.

**Figure 4.**
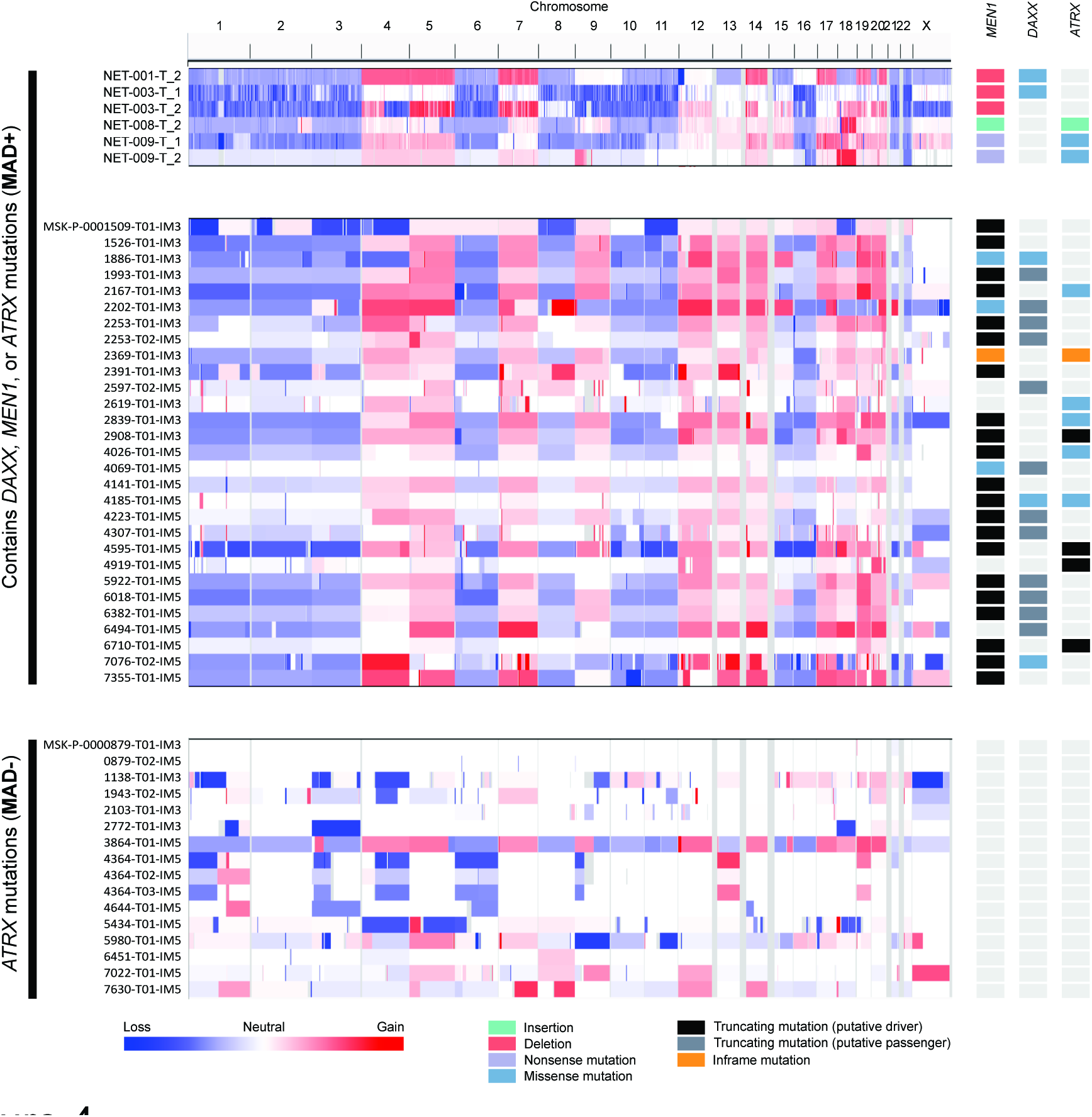
Relative copy-number profiles for pancreatic neuroendocrine tumours from the discovery as well as the publicly available AACR GENIE PNETs cohort. Samples are divided based on the presence mutations on *MEN1, ATRX* or *DAXX* (MAD+), or the complete absence of mutations on any of these genes (MAD-).

To test whether MAD mutations arose prior to LOH and genome doubling events, we adapted our molecular timing analysis for the GENIE cohort. Due to the lack of absolute copy-number values, we used pathologist estimates of tumour purity (+/- 0.15) paired with the simplest copy-number models that best explained the observed allelic fractions for all mutations and maintained constraints imposed by copy-number status (i.e. *DAXX* and *MEN1* have the same copy-state) (Supp. Table 24). Of the MAD+ GENIE PNETs, 6/29 had to be removed due to estimated purities below 30%. The remaining 23 PNETs showed a strong tendency to adopt a copy-number model that reinforces the idea that *MAD* mutations occurred prior to LOH and genome doubling events (Fig. 3). As tumour purities are rough estimates by pathologists, we have some uncertainty regarding the most likely copy-number model. For instance, outliers with a cancer chromosome fraction greater than 1.2 are biologically impossible as this model implies the mutation is on 120% of all tumor-specific chromosomes. However, the data still accurately place the acquisition of mutations before and after LOH and doubling events.

Overall, we observed a significant enrichment of *MEN1* and *DAXX* mutations prior to LOH and genome doubling events (Bonferroni adjusted p-values: *MEN1* =0.00029, *DAXX* =0.00011, binomial test) when using a stringent cutoff of 0.85 cancer chromosome fraction. *ATRX* reached significant enrichment at a cancer chromosome fraction cutoff of 0.63, which may reflect difficulty in estimating copy-state for chromosome X. Assuming that all pathologist estimates of tumour cellularity are correct, we observe 35/39 MAD+ PNET samples follow our molecular timing model (4/4 discovery cohort, 10/12 validation cohort, 21/23 GENIE cohort) (p.value= 3.353e-07, binomial test). Therefore, we conclude that acquisition of *MEN1* and *DAXX/ATRX* mutations are initiating events that lead to a single catastrophic LOH-event, frequently followed by whole genome doubling.

### Potential tumor-suppressor genes in LOH regions

We next sought to uncover whether there was a copy-number dependent effect on expression as a result of the PNET CN-LOH signature. We compared the overall expression of genes within consistent heterozygous and LOH ranges for both our PNET and GINET samples, as well as pancreatic samples from the Genotype-Tissue Expression project (GTEx). Expression within heterozygous and LOH regions did not differ between PNET, GINET, or normal-pancreatic samples (Supp. Fig. 9a). Additionally, we observed no significant changes in expression between LOH and heterozygous gene-expressions within the same sample. However, all GINET samples displayed a 1.32 fold lower median expression of heterozygous genes in comparison to LOH genes (Supp. Fig. 9b).

Loss of heterozygosity may be functioning as a second-hit mechanism that could potentially uncover tumor suppressor genes (TSG). To explore this, we established a normal level of expression by generating a probability density function for each of the 31 tissue types in GTEx. Potential TSGs within conserved LOH regions were nominated based on the criteria that all PNET samples within our discovery cohort had less than a 5% probability of being expressed at a normal or higher than normal level expression across 95% of all tissues. Using this method, we nominated 43 potential TSGs, 18 of which were found at lower probabilities in PNETs (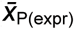 < 0.05) compared to GINETs (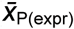 > 0.10) (Table 3). However, one key limitation of this approach is that this is not a definitive indication of loss-of-function in TSGs, as we can not accurately measure ablation of expression while there is normal tissue contamination. More formally, this is a measure of gene expression that is lower than what we would expect in healthy tissue driven predominantly by a low expression in tumor content.

**Table 3.**
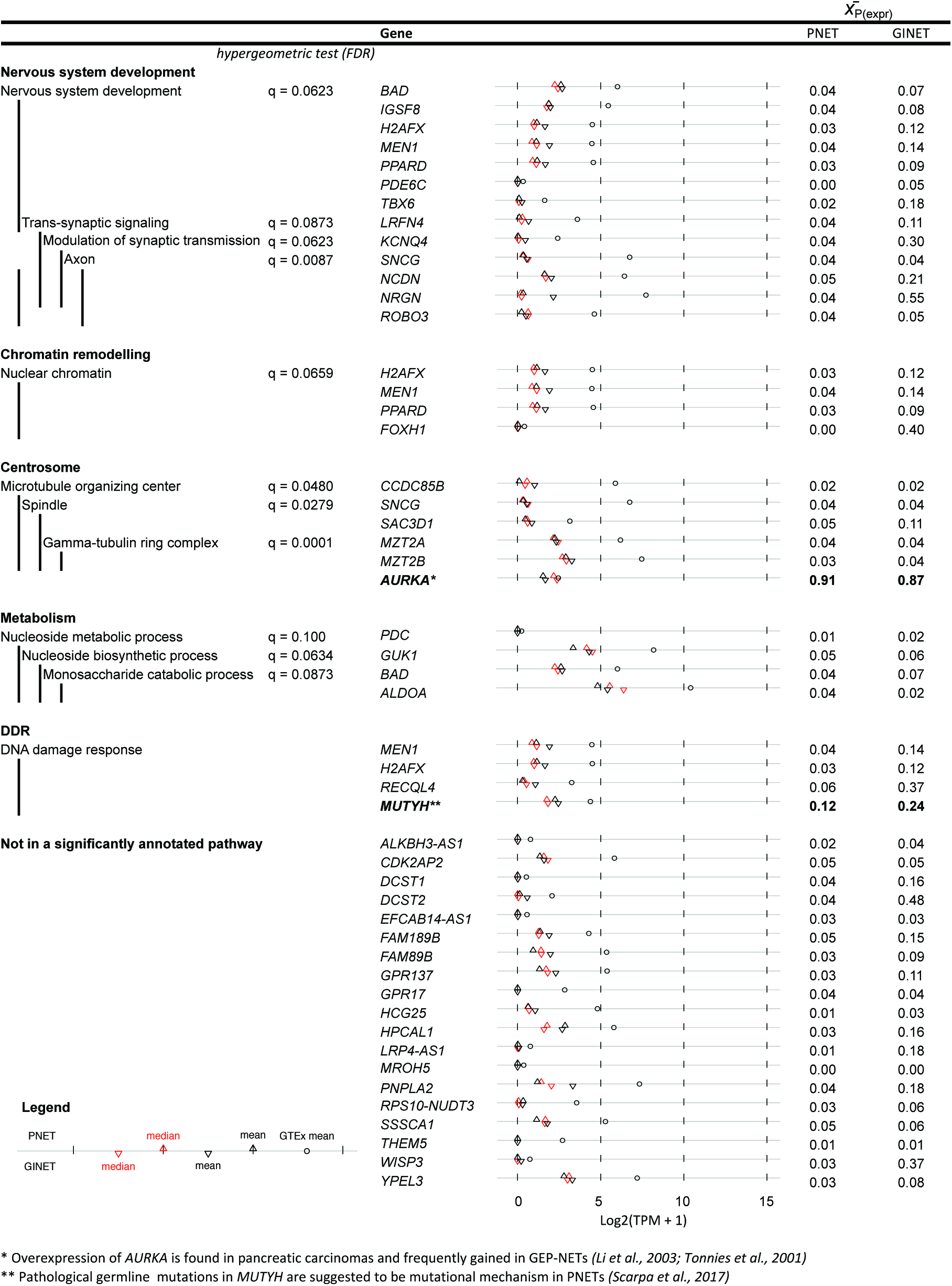
The lowest expressed LOH genes in PNET and GINET samples of the NETseq discovery cohort in comparison to the basal expression of those genes in samples covering all GTEx tissue types. Genes were selected if the mean percentile of PNET genes were below the 5^th^ percentile 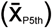 of this comparative GTEx distribution. An over-representation analysis performed using a hypergeometric test was done using ConsensusPathDB.

An over-representation analysis of these 43 lowly expressed PNET genes showed significant enrichment for molecular functions and components such as nervous system development (13/43 genes), chromatin remodelling (4/43 genes), microtubule organizing center (5/43 genes), and metabolism (4/43). The remainder of the genes were not significantly enriched (q > 0.1) in any ontology geneset other than the non-specific “intracellular non-membrane-bounded organelle” (Supp. Table 25). Additionally, there was no geneset that favored loss of PNET genes over GINET genes.

Of the genes composing the neuronal component, 6 are enriched in trans-synaptic signaling and axonal components: *SNCG*, *NRGN, NCDN, KCNQ4, ROBO3, and LRFN4*. The biological significance of these genes and ontology in neuroendocrine cells is currently unknown but it may be related to the non-functional component of PNETs via a diminished secretory mechanisms of antigens. Additionally, there appeared to be no preferential loss of these genes in LOH regions of PNETs compared to GINETs; suggesting a tissue non-specific method of dimished antigen secretion.

Another striking ontology was the loss of genes enriched in the microtubule organizing center (MTOC), a major component of centrosomes and required for normal amphitelic attachment and chromosome segregation during cell division^26^. In agreement with previous PNET^13^ and human pancreatic cancer studies^27^, we have also identified a recurrent copy-number gain of chromosome 20q13, a region harboring *AURKA*, paired with overexpression relative to all GTEx tissues (PNET: 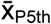 = 0.91; GINET: 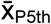 = 0.87). Centrosome amplification is known to be associated with merotelic attachments of microtubules to kineotchores, resulting in lagging chromosomes and chromosomal instability in bipolar cell divisions^28^. Meanwhile, cells with centrosome depletion are still able through spindle assembly checkpoint evasion, but they do not always lead to increased chromosome missegregation^29^. It is unclear from these results whether centrosomes are lost or gained in PNETs, however, the loss and gain of essential MTOC genes suggests perturbation of the centrosome and a potential mechanism of chromosomal instability.

The DNA damage response gene set did not prove to be significantly enriched in our analysis. However, we have established functional relevance based on the literature and relevance to NET development. In agreement with the molecular profiles of our PNETs, the biallelic inactivation of the TSG *MEN1* through nonsense and truncating mutations paired with LOH resulted in decreased expression compared to all other GTEx tissues. The loss of this tumor-suppressor has pleiotropic effects, ranging from histone modification, DNA methylation, to affecting homologous recombination (HR) and non-homologous end-joining (NHEJ) DNA repair. Through this analysis, we have also identified a loss of expression for *H2AXF*, or what is referred to as the the “histone guardian of the genome”^30^. The depletion of this gene has been shown to be correlated with poor prognosis and increased genomic instability, most likely through the response of H2AX double-stranded breaks^30,31^. Finally, we also nominated loss of cancer-syndrome associated genes *RECQL4* and to a lesser extent, *MUTYH*, both of which did not meet our percentile cutoff but were associated with large discrepancies between PNET and GINET percentiles. Mutations in the base-excision repair gene *MUTYH* has recently been shown as a feature seen in PNETs but absent in pancreatic adenocarcinomas^24^. Meanwhile, RecQ Like-Helicase 4 works in a complex to dismantle aberrant DNA structures of repetitive genomic regions such as G-quadruplexes that stall replication forks. Loss of this gene is associated with the cancer Rothmund-Thompson and is known to be associated with mosaic aneuploidies, isochromosomes, and mis-segregations^32^.

The remaining genes nominated by our analysis were not enriched in any ontology or pathway geneset, nor was there any literature relating their function back towards chromosomal instability or neuroendocrine biology. Additionally, while *ATRX* and *DAXX* also show biallelic inactivation within our exome sequencing analysis, we did not observe loss of expression of these genes in PNET or GINET samples. This is most likely due to a missense mutation rendering the gene non-functional but still transcribed, rather than a truncating or frameshift mutation as seen in *MEN1*.

## Discussion

Previous studies have shown that PNETs with a mutation in *DAXX/ATRX* are subject to chromosomal instability and describe a more aggressive phenotype^8^. Of this subgroup, the presence of a *MEN1* mutation is associated with longer overall survival^4^. In our study, we found that this subset of aggressive PNETs is further characterized by CN-LOH that afflicts the majority of the genome. The enrichment of the copy-neutral LOH afflicting large portions of PNET genomes showed a trend towards increased metastasis in our validation cohort, consistent with copy ratios of the same chromosomes seen in publicly available CGH datasets. Thus, we hypothesize that metastatic PNETs are defined by this copy-neutral loss of heterozygosity signature; likely triggered by loss of *DAXX/ATRX* and subsequently facilitated by biallelic loss of *MEN1*.

Despite this extensive LOH feature being previously described in SNP allelotyping and microsatellite analysis studies^22,33,34^, we show here that this signature is highly defined within our discovery and validation cohort and likely makes up a subgroup of metastatic PNETs. Previous allelotyping studies using microsatellite loci also demonstrated this widespread LOH pattern, however, the severity of this signature was obscured by the lack of parallel copy-number profiles^33,34^. One study paired these allelotype-profiles with flow-cytometry inferred ploidy; 5 of the 7 samples that presented with a high fractional allelic loss (FAL ~ 0.55) were also aneuploid tumours. These 5 tumours maintained a similar pattern of LOH status as those observed in our discovery and validation cohort, suggesting that they may occur in a copy-neutral state^33^.

A more recent study using SNP allelotyping presented a copy-number/LOH profile that is discordant to those described in our study^22^. While we present a distinct CN-LOH signature that appears to be highly retained within our discovery and validation cohort, Nagano and colleagues only observe CN-LOH in chromosome 3 for a single PNET. While this study describes a copy-number profile that is highly similar to those observed within our discovery/validation cohort, it also suggests that losses of chromosomes 1, 2, 3, 6, 8, 10, 11, 15, 16, 21, and 22 are recurrently paired with gains of the remaining chromosomes in 5 of the 9 cases that display high chromosomal instability. The copy-profile associated with these 5 tumours are slight outliers when compared to other array-CGH, exome-sequencing, and shallow-WGS studies presented in our paper. While these cases may be originating from a heterogeneous subpopulation of PNETs as the authors suggest, we propose that scaling of the copy-state may be affected by normalizing a tumour genome that is half-diploid and half-tetraploid to a diploid genome; possibly resulting in the incorrect centering of these copy-number profiles to an overall ploidy of 2 instead of a ploidy of 3.

The biallelic loss of *ATRX* /*DAXX* paired with *MEN1*, as well as PNET-specific loss of expression for *H2AFX* and *RECQL4* all share a common fundamental roles in DNA-damage repair and epigenetic progression of cancer. The tri-protein MRN complex (MRE-11, RAD50, and NBS1) is involved in HR and NHEJ, as well as responding to replication fork arrest^35^. In mice, knockout or haploinsufficiency of H2AX is related to decreased MRN accumulation at sites of double-stranded breaks (DSBs), decreased efficacy of HR and NHEJ-mediated DNA repair, as well as an increase in chromosomal aberrations^30,31,35-37^. Through knockout studies, all other gene products for *MEN1*, *ATRX/DAXX*, and *RECQL4*, also show deficiencies in DSB repair by impairing HR-mediated DNA repairs and impediment rectifying stalled replication forks at aberrant DNA structures^38^^-^^40^. Phosphorylation of the histone variant H2AX to γH2AX

Related to the aforementioned mechanisms of impaired DNA damage response, these genes are proposed to promote chromosomal instability via epigenetic roles. the gene-product of *MEN1*, menin, is involved in recruiting the H3K4me3 histone methyltransferase mixed-lineage leukemia (MLL) complex and functions as a potent tumor suppressor in pancreatic islet cells^41^. Menin exerts a pleiotropic effect by interacting with over 2,000 gene promoters^42^, several of which are involved in HR response to DSBs^38^, others of which are involved in genome-wide hypermethylation leading to increased activity of Wnt/β-catenin signalling^43,44^ and down-regulation of p27 expression^45^. Meanwhile, the DAXX/ATRX complex is linked to the deposition of histone variant H3.3 at heterochromatic regions of the genome; specifically, re-establishing H3.3-containing nucleosomes at intrinsically repetitive regions that form G-quadruplex structures and stall replication forks^41,46^.

An alternative lengthening telomere phenotype induced by the loss of *ATRX* and/or *DAXX* can also promote chromosomal instability in late-stage PNETs^8^. Therefore, we hypothesized that these are likely initiating mutations leading to subsequent aneuploidy seen in PNETs. This acquired LOH state may be essential for PNET progression by revealing underlying mutations or allele-specific DNA methylation patterns, the latter possibly due to tissue-specific or parental imprinting.

Due to the well-conserved nature of the CN-LOH signature and the strong association to be preceded by mutations in both *MEN1* and *DAXX/ATRX*, we rationalize that these mutations are likely to be driver mutations that initiates LOH events. As a means to stabilize an unstable hemizygous genome, PNETs would adopt a genome-doubled state that is necessary for the metastatic phenotype and increased fitness of the cells. The resulting expression from homozygous genes displays a consistent lack of expression for genes that play a role in centrosome formation and DNA damage response; factors that would contribute to further chromosomal instability and a more dynamic karyotype. Likewise, there must be a network of components present in the chromosomes that always retain heterozygosity that are essential for PNET survival. Thus, a better understanding of the functional implications of regions of retained zygosity on the background of this CN-LOH signature may suggest therapeutic targets to treat this disease and uncover selective mechanisms underlying aneuploidy in cancer.

## Methods

### Tissue acquisition

Our discovery cohort originated from 7 patients enrolled in the NET-SEQ study at the Princess Margaret Cancer Centre. Eligible patient had histological or cytological diagnosis of well-differentiated neuroendocrine tumours (NETs) or well-differentiated pancreatic neuroendocrine tumours (PNETs). Our validation cohort comprised of 38 NET samples provided by the Ontario Tumour Bank, which is supported by the Ontario Institute for Cancer Research through funding provided by the Government of Ontario. Three sample types were processed: buffy coat blood cells, formalin-fixed paraffin embedded (FFPE) tissues at time of diagnosis, and fresh-frozen core needle biopsies.

### Genomic characterization

DNA exome, RNA-sequencing was performed on the discovery cohort, resulting in 250x coverage in tumours, 50x coverage in normals, and ~80,000,000 reads for RNA-sequencing. Sequence data was aligned the reference sequence build hg19. Variant detection in exome data was performed using MuTect^47^ and HaplotypeCaller^48^, while copy number data was called using VarScan2^49^ and Sequenza^50^. Loss-of-heterozygosity data was inferred from both DNA and RNA data by determining purity-adjusted allelic fractions. Gene-wise transcript abundances were quantified using the Cufflinks suite of tools. Shallow whole-genome sequencing was performed on fresh-frozen samples of the validation cohort, resulting in 0.4X coverage in tumours. Copy-number profiles were obtained by decomposing the aggregate coverage across the genome using a mixed-Gaussian distribution model into individual copy-state distributions. Loss of heterozygosity was estimated by comparing the number of heterozygous variants (defined as an allelic fraction not 0 or 1) within a 500kB bin (~26 heterozygous variants per bin) to a reference empirical cumulative density function composed of all 500kB bins across all tumour genomes. By selecting the most prevalent percentile across each copy-number segment, we termed a chromosomal segment as either LOH or heterozygous based on a model trained on pathologist-estimated stromal content. To validate these copy-number calls, we paired this analysis with fluorescence in-situ hybridization on complementary formalin-fixed paraffin embedded tissues.

### Meta-analysis of published datasets

Whole-exome sequencing of the BON-1 and QGP-1 PNET cell line from VanDamme and colleagues ^14^ was re-analyzed and LOH segments were called based on allelic fractions (European Nucleotide Archive study ID: <u>PRJEB8223</u>). Allele specific copy number data for a subset of the TCGA PanCancer cohort were downloaded from the supplementary data files accompanying initial report of the ABSOLUTE algorithm ^16^. Loss-of-heterozygosity and copy-states for each copy-number segment were taken directly from these published results and were used to estimate the fraction of the genome afflicted by the LOH state. Copy number profiles derived from comparative genomic hybridization microarray data were obtained from data tables described in six publications^11-13,17-22^ and transcribed into genomic coordinates by mapping to cytobands using the UCSC Table Browser hg19 cytoBandIdeo file (http://hgdownload.cse.ucsc.edu/goldenPath/hg19/database/cytoBandIdeo.txt.gz). Each copy ratio segment was assigned a value corresponding to the copy-status. Jaccard index values were calculated to measure the asymmetric binary concordance between any two copy-number profiles.

### Detection of tumor-suppressor genes

Gene-wise transcript abundances were reprocessed for GTEx RNA-seq data using the same STAR (v2.4.2) two-pass method paired with Cufflinks. Empirical cumulative density functions (ECDF) were generated for each gene across all samples within each of the 31 tissue types. For each sample with RNA-seq data in our discovery and validation cohort, each gene within the LOH regions was projected onto the gene-level ECDF for each tissue type to identify the corresponding percentile. For each sample, the upper-end of all percentiles across all of the 31 GTEx tissue types was taken as a representative percentile to estimate how expression compares against GTEx. Potential TSGs were nominated based on the average percentile for PNETs being less than 0.05 and the average percentile of GINETs being larger than than 0.1. An over-representation analysis was done on these genes using ConsensusPathDB (http://cpdb.molgen.mpg.de/CPDB), which executed a hypergeometric test paired with a false discovery rate multiple hypothesis testing adjustment.

### Molecular Timing in Project GENIE

Copy-number profiles and mutational data of PNETs from AACRs project GENIE (v1.0.1) were downloaded from Sage Synapse (https://www.synapse.org/; synapse IDs: syn7851250, syn7851253, and syn7851246). In total, 43 PNET samples had both copy-number information and mutational information. The molecular timing of these samples was determined by estimating the tumour purity required for every possible copy-number profile to generate the observed tumour purity for all somatic mutations. The simplest copy-number profile that fit the constraints of pathologist purity +/- 0.15 and copy-number constraints imposed by the relative copy-states of the somatic mutations was used to infer molecular timing of the disease.

Additional methods and detailed version and parameter information are available in the Supplementary Methods.

## Acknowledgements

This project was funded by grants from the Cancer Research Society and the Carcinoid NeuroEndocrine Tumour Society Canada (#19341, TJP) and the Princess Margaret Cancer Centre Neuroendocrine Tumor Research Fund (LLS). Infrastructure support was provided by the Princess Margaret Cancer Foundation; Canada Foundation for Innovation, Leaders Opportunity Fund, CFI 340 #32383; and Ontario Ministry of Research and Innovation, Ontario Research Fund Small Infrastructure Program (TJP). RQ is supported by a Medical Biophysics Excellence Ontario Student Opportunity Trust Fund Award from the Princess Margaret Cancer Centre, the Province of Ontario, and the University of Toronto. AS is supported by a Conquer Cancer Foundation ASCO Young Investigator Award. We thank the staff of the Princess Margaret Genomics Centre (www.pmgenomics.ca, Neil Winegarden, Julissa Tsao, Nick Khuu, and Gurbaksh Basi) and the Bioinformatics and High-Performance Computing Core (Carl Virtanen, Zhibin Lu, and Natalie Stickle) for their expertise in generating the sequencing and microarray data used in this study. We also thank Michael F. Berger of the Memorial Sloan Kettering Cancer Center for facilitating pathology estimates for samples within the AACR GENIE cohort. Biological materials were provided by the Ontario Tumour Bank, which is funded by the Ontario Institute for Cancer Research.

